# Integrated Database of Force-Field Parameters, Experimental Measurements and Molecular Dynamics Simulations

**DOI:** 10.1101/2024.12.03.626554

**Authors:** Pavel Banáš, Vojtěch Mlýnský, David Číž, Radek Furmánek, Nestor Pilat, Viktoria Pauw, Stephan Hachinger, Jiří Šponer, Jan Martinovič, Michal Otyepka

**Affiliations:** IT4Innovations, VSB – Technical University of Ostrava, 17. listopadu 2172/15, 708 00 Ostrava-Poruba, Czech Republic; Czech Advanced Technology and Research Institute, CATRIN, Křížkovského 511/8, 779 00 Olomouc, Czech Republic; Institute of Biophysics of the Czech Academy of Sciences, Královopolská 135, 612 00 Brno, Czech Republic; Leibniz-Rechenzentrum(LRZ), Boltzmannstr. 1, 85471 Garching bei München, Germany

## Abstract

Molecular Dynamic (MD) simulation is a vital theoretical tool for exploring nucleic acids (RNA, DNA), proteins and other (bio)molecular systems, generating vast amounts of data daily. Efficient storage and possible reuse of this data is a persistent challenge. Here, we introduce IDA (Integrated DAtabase of force fields and datasets from experiments and MD simulations), an innovative database scheme for datasets from various types of MD simulations. IDA supports outputs from different MD approaches, i.e., standard MD simulations, importance sampling techniques, simulated annealing, and other enhanced sampling methods including replica-exchange simulations. IDA also houses a collection of molecule-specific force fields (FFs) and experimental datasets. Uploaded MD outputs, FFs, and experimental data are integrated in a standardized format, allowing efficient data mining and extraction of valuable insights from the extensive data generated by diverse MD simulations. With the data and metadata holdings of IDA, and the prospective assignment of persistent identifiers, our work aims to make key steps towards making MD data FAIR (findable, accessible, interoperable, reusable).

## BACKGROUND & SUMMARY

Classical molecular dynamics (MD) simulations serve as a vital theoretical tool used for exploring the structure, dynamics, and function of a wide range of molecular systems, from small molecules to biomacromolecules and nanomaterials.^1-4^ These simulations integrate Newton’s equations of motion over time using short integration steps (on a femtosecond timescale), with forces calculated using an empirical potential function called a force field (FF). Leveraging high-performance computing (HPC), MD simulations can generate vast amounts of data, necessitating substantial processing and storage for thorough analysis. Annually, approximately 40,000 scientific papers on MD are published, a number that has doubled over the past decade. During this period, the typical simulation time has extended from tens of nanoseconds to tens of microseconds. Consequently, the processing and analysis of MD simulation data have become significant bottlenecks in the field. Although both HPC and MD simulation methods have witnessed significant progress over the past decades, considerably less attention has been paid to the methods and technologies for data processing and analysis.

The first challenge associated with MD simulations-based data is their efficient storage not only due to large amount of data (from GB to TB per one MD simulation) but also due to efficient access to data for their processing and analysis. Huge amount of output data from simulations hampers their upload to public repositories and thus, if so, only portion (or truncated) datasets are eventually uploaded. Both storage and reusability of MD datasets are further complicated because datasets may vary in formats due to MD engine (e.g., AMBER, GROMACS, OPEN-MM, Tinker-HP, etc.) and structure due to simulation setup (e.g., standard single trajectory, multiple replica exchange trajectories, biased trajectory, etc.). Hence, the variety of data sets between different MD simulation methods significantly limits interoperability of stored data which must be merged, classified, and stored together in standardizable format to allow efficient reusability.

Unlike structural data, which are publicly available for decades and efficiently organized in the Protein Data Bank (https://www.rcsb.org/),^5^ there is a visible gap in publicly available databases integrating FF parameters, a diverse range of MD simulation data, together with experimental data corresponding to the simulated system. Initial community efforts have focused on generating trajectory databases specifically for protein simulations, as exemplified by initiatives such as Dynameomics^6^, MoDEL^7^, and Dynasome^8^. They contain collections of hundreds to thousands very short protein simulations and often required specific software tools for the analysis. Cheatham and coworkers introduced an infrastructure for managing and sharing simulation data, iBIOMES^9^, based on the iRODS framework (https://irods.org). This system provides a straightforward mechanism for data deposition and allows for the registration of data into the system without relocating it from its original storage location. More recently, Orozco group introduced two database platforms, i.e., BIGNaSim^10^ and BioExcel-CV19^11^. The BIGNASim platform provides access to trajectories from MD simulations of DNA molecules, including pre-computed analyses and other analyses performed on the database. Additionally, the BioExcel-CV19 database offers web-based access to atomistic MD trajectories for macromolecules related to COVID-19. The database can accommodate trajectories from different MD engines and different kinds of MD simulations, which are upon upload converted into an in-house binary format. The BioExcel-CV19 also contains sets of pre-computed analyses and it is ready for future development under Molecular Dynamics Data Bank (MDDB) project.^12^ The NMRlipids Databank is a community-driven, openly accessible catalogue that houses MD simulations of lipid membranes.^13^ It successfully integrates experimental data from NMR and X-ray scattering techniques. The databank facilitates user-defined automatic evaluations of trajectories, allowing simulations to be ranked based on their FF performance against available experimental benchmarks.^13, 14^ All those past efforts indicate progress in building comprehensive MD simulation databases, but they primary focus in developing tools for accessing and analyzing individual simulations. However, none of these existing database formats enable capabilities for data mining and extracting value from the vast amounts of data generated by all these (diverse) stored simulations together. Data mining on a sufficiently robust simulation database can, e.g., offer valuable insights into the performance and accuracy of the employed FFs, the effects of particular simulation settings, etc.

Another challenge relevant to the further advancement of MD simulations is the integration of MD data with other relevant data sources, including FFs and experimental data. The interconnection of these data sets can be utilized to assess the relevance of MD simulations, test new simulation methods, and develop advanced analytics tools. These advancements can be significantly accelerated by machine learning (ML) tools, which require data that is accurate, consistent, well-structured, and relevant. Moreover, the seamless integration of diverse data sources will enhance the reliability and predictive power of MD simulations, fostering innovation in fields such as drug discovery, materials science, and biotechnology. Continued efforts to develop standardized data formats and interoperability frameworks will be crucial in achieving these goals.

In this work, we propose a novel standardized database IDA (Integrated DAtabase of FFs and datasets from experiments and MD simulations) for storing datasets from diverse types of MD simulations. This includes datasets from standard MD simulations as well as those from advanced techniques such as importance sampling techniques, simulated annealing, and other enhanced sampling methods including replica-exchange simulations. The IDA also stores experimental datasets, primarily from NMR experiments, along with metadata from user-specified analyses. These analyses are calculated, visualized, and most importantly, stored for immediate visualization upon subsequent requests by different users. The database adheres to Findability, Accessibility, Interoperability, and Reusability (FAIR) principles^15^. Currently IDA contains precalculated datasets from MD simulations of small RNA motifs and benchmark experimental data for the comparison. The database is also housing a collection of molecule-specific FFs (AMBER^16^ type formats), i.e., FF parameters for each type of biomolecule (RNA, DNA, protein etc.), ions and solvents. These molecule-specific FFs are stored as separated downloadable files.

## METHODS

### General IDA structure

IDA integrates four major interrelated databases containing FFs, experimental data, MD simulations and analyses derived from MD simulations. In addition, IDA includes a user database for assigning roles and each database contains metadata (Figure 1). A record in the user database is created upon user registration and validation of their email address. Besides IDA Administrator, four levels of user rights are assigned: (i) guest, (ii) user, (iii) MD Expert, and (iv) IDA Expert, with roles and rights listed in Table 1. Guests has completely free access to the IDA database. User, MD and IDA Experts are required to register (via email account) and the registration is approved by the IDA Administrator. Upon completion, records in the databases of FFs, experimental data, and MD simulations are locked and automatically annotated with a PID, upload_user_name, and upload_timestamp. The detailed structure of IDA is shown in Figure 2.

**Table 1:** List of different IDA users with their roles and rights.

| User | Roles and Rights | Access |
| --- | --- | --- |
| Guest | reads data | Free |
| User | reads data, executes analyses | Registration |
| MD Expert | same as User, uploads MD simulations | Registration |
| IDA Expert | same as MD expert, uploads experimental data and FF parameters | Registration |
| IDA Administrator | assigns user roles | Registration |

**Figure 1:**
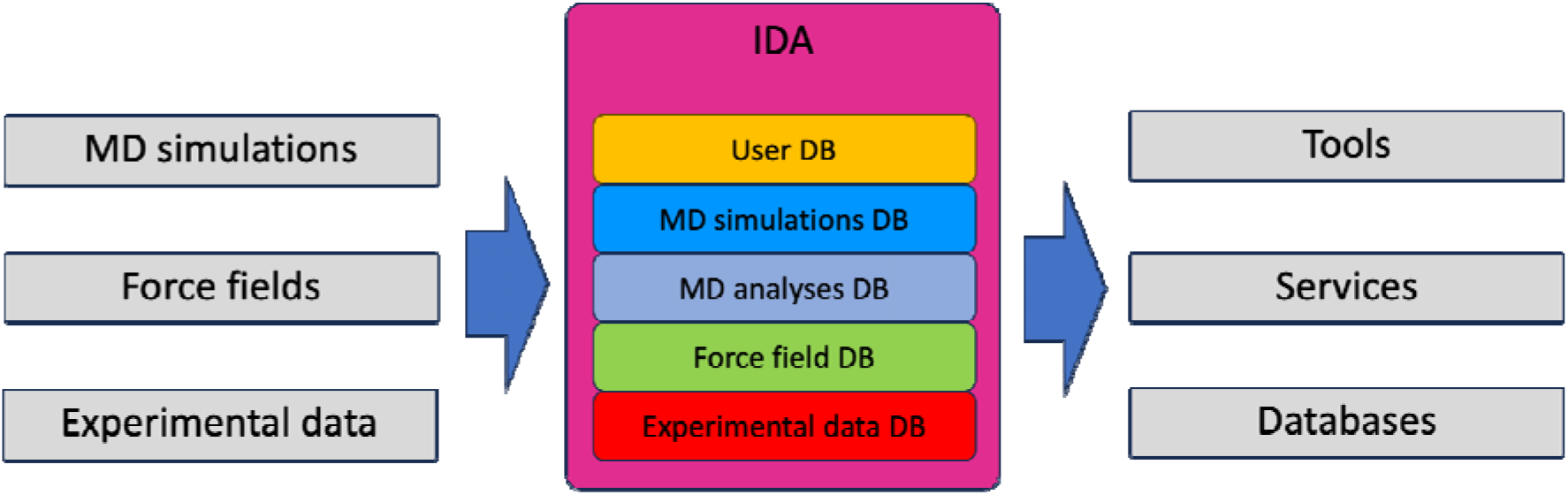
Scheme shows data sources and their integration in the IDA database. The IDA structure is composed by five interrelated databases, which are highlighted by different colors.

**Figure 2:**
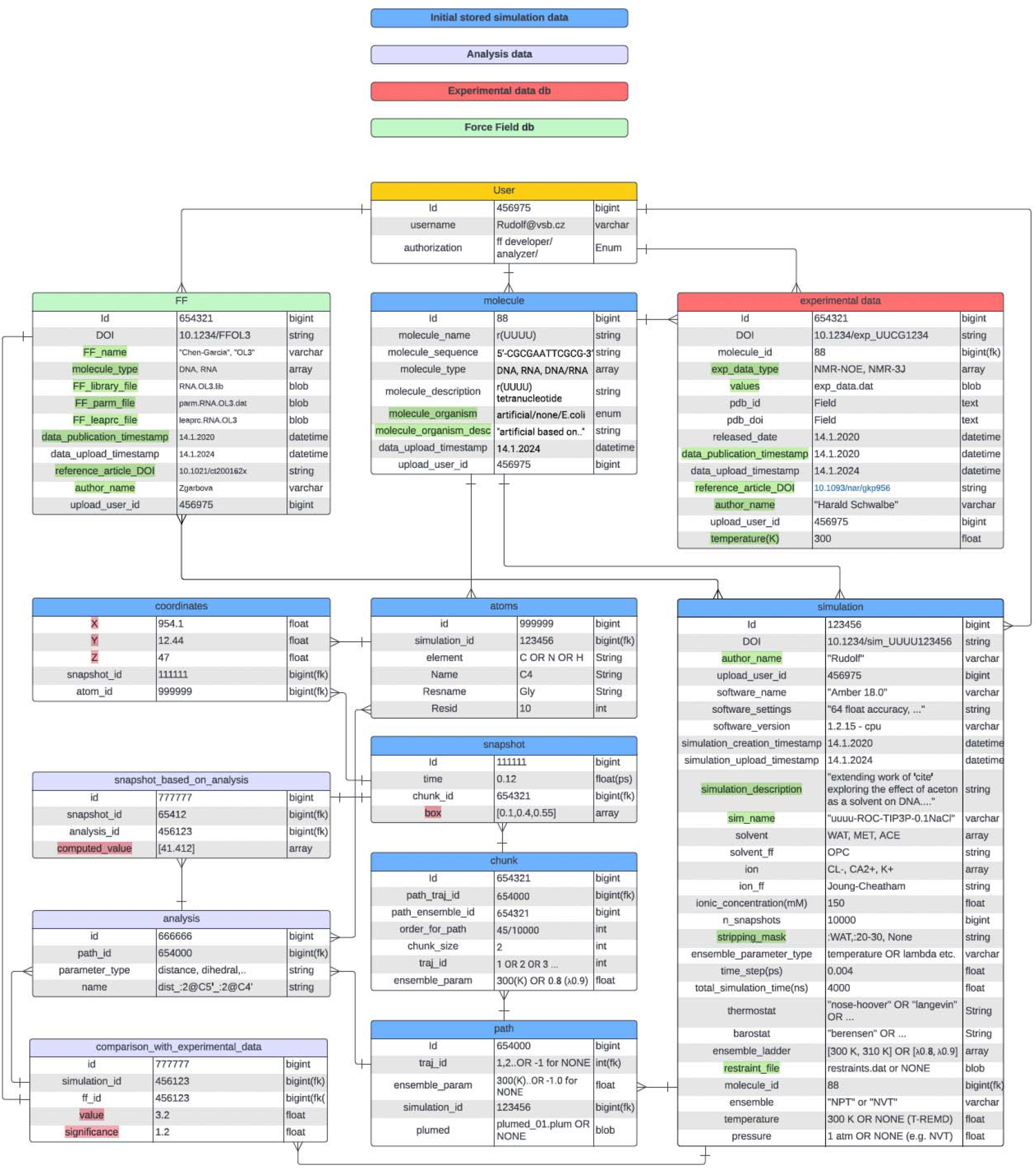
Detailed scheme of the IDA database. The items that are required to be added manually by user are highlighted in green. The variables and files containing data are highlighted in red, the other items are considered as metadata.

### Database of FFs

The FF records are entered only by IDA Experts, who are acknowledged for their expertise in database structure and its purpose. The users must provide the following metadata: FF name (FF_name), author name (author_name), date of publication of this FF (data_publication_timestamp), and reference to the original research article in the form DOI (reference_DOI). The user selects one of the types of the molecule in the FF, e.g., RNA, DNA, protein, water etc. The list of available molecule types can be extended only by IDA Administrator. In addition to the metadata, users must upload three files containing FF data; i) AMBER FF object file format library file (FF_library_file), ii) AMBER FF parameter file (FF_parm_file), and iii) source file for LeaP AMBER program (FF_leaprc_file). Detailed information about these file formats can be found at AMBER File Formats web page (https://ambermd.org/FileFormats.php). Currently, IDA supports AMBER file formats. It also verifies that the uploaded FF corresponds to the selected molecular type based on the uploaded files.

### Database of Experimental Data

Experimental data records can only be entered by IDA Experts. Users must provide the following metadata: the author of the experimental data (author_name), the publication date (data_publication_timestamp), a reference to the original research article in the form of a DOI (reference_DOI), and the temperature at which the experimental data were obtained (temperature). Additionally, users must specify the type of measurement from a predefined list (exp_data_type), which can be expanded to include new experimental data types upon approval by the IDA Administrator. Examples of measurement types include NMR-3J couplings, NMR-NOE, NMR-unobserved NOE, etc. Users are also required to associate the experimental data with a specific molecule (Figure 2). The system automatically links the MD simulations to the corresponding experimental data. Experimental data are uploaded as files (exp_data_file) in formats compatible with the selected data type, such as NMR-STAR or NEF file formats as used in PDB database (https://www.rcsb.org;^5^ see, e.g., Refs. ^17, 18^). The IDA system verifies that the uploaded file format matches the selected type of experimental data.

### Molecule Records

The molecule records are part of MD simulation database. These records may be entered by MD Experts and IDA Experts, i.e., skilled domain experts. The molecule records are connected to the experimental data and MD simulations data sets, thus pinning the MD simulations to the corresponding experimental data. Once the MD simulation is uploaded (see below), the IDA identify the biomolecule in the system topology file and associate the simulation with the corresponding molecule record (if it already exists) or create a new molecule record. In such cases, users are required to add information regarding the organism(s) from which the molecule was originally derived (molecule_organism), if applicable, and a concise description of the molecule (molecular_organism_desc, e.g., *E. Coli*). All other information are automatically parsed from the topology file.

### MD Simulation Database

The records in the MD simulation database may be entered by MD Experts and IDA Experts. Authorized users enter AMBER trajectory file(s), topology file(s) and output logfile(s). The data and most of metadata stored in the database will be extracted automatically from these files, however, the database will not save the files themselves to ensure future compatibility with other file formats. The system verifies the completeness of the files on input, e.g., if each trajectory is complemented by the corresponding output logfile etc., and flags any discrepancies in the data. Beside the information parsed from trajectory, topology and output logfiles, users are asked to enter following additional metadata; author of the simulation (author_name), name of the MD simulation (sim_name), restraint file containing information about applied restrains including gHBfix (restraint_file - if applicable), and stripping mask if applicable (stripping_mask). The stripping mask will be included when the simulation molecular system as uploaded into the database was stripped out compared to the complete system used for generation of the simulation, typically water molecules, ions etc. The stripping mask (in the same format as used by CPPTRAJ AMBER tool)^19^ thus identifies what atoms were stripped out. Although the IDA is currently open to outputs only from AMBER simulation package^16^, it is designed to accommodate outputs from different MD engines, e.g., GROMACS^20^, OPEN-MM^21^, TINKER-HP^22^, etc.

The other items in simulation record will be automatically parsed from the topology and output logfiles. Namely, name of the MD engine software (software_name), setting of the MD engine precision (software_settings), software version (software_version), time and date of the simulation creation (simulation_creation_timestamp), number of snapshots in the simulation (n_snapshots), identification of the ensemble parameter in replica exchange simulations (if applicable – ensemble_parameter_type), integration time step (time_step), total simulation time (total_simulation_time), identification of barostat (if applicable), identification of thermostat (if applicable), values of the ensemble parameters in replica exchange simulation (if applicable – ensemble_ladder), thermodynamic ensemble, temperature (if temperature is not the ensemble parameter, such as in T-REMD), and pressure (if applicable) are taken from output log files. The information about solvent, ions and ionic concentration are taken from topology file. Based on information from topology file, the simulation is linked to the partial molecule (see above) and particular FF for each type of molecule in the molecular system, so that the IDA compares the FF parameters in the topology file with those in FF database and automatically identifies and validates the FF.

The atom records are taken from topology file and include atomic metadata information, i.e., atom name, atom id, identification of element, name of the residue, where the atom belongs to, and residue id. The trajectory files are parsed to the path, chunks, snapshots, and coordinates records, so that both trajectories and ensembles, i.e. both complemented sets of simulation paths are represented explicitly listed as path item in the database. In case of standard MD simulation, only one path is present. The paths might involve trajectory id (simulation_id) and value of ensemble parameter (ensemble_param). If path is trajectory, the simulation_id contains number of trajectory (following time evolution of the coordinates) and ensemble_param is set to none, while in case of ensemble, the simulation_id is set to none and ensemble_param involves ensemble parameter value. Thus, the database can clearly determine, whether path corresponds to trajectory or ensemble. Each path is associated to chunks that belongs to that particular path. Every chunk has set both simulation_id and ensemble_param values determining to what trajectory and ensemble the chunk belongs, and corresponding IDs of ensemble path and trajectory path. In addition, chunk contains the information about position of the chunk in its ensemble and trajectory paths (order_for_path) and number of snapshots in the chunk (chunk_size). Every chunk is associated to its snapshots containing information about simulation time (time) of the snapshot, and size of the simulation box (if applicable - box). Every snapshot is associated to its coordinates containing Cartesian coordinates and are linked to the particular atoms (Figure 2).

## DATA RECORDS

### MD Simulation datasets

Classical all-atom MD simulations describe the behavior of a molecular system, e.g., a biomolecular solute like RNA, DNA, protein surrounded by counter ions and explicit water molecules over time. A typical MD simulation generates a continuous series of atomic coordinates at regular time intervals, collectively referred to as an MD trajectory. The trajectory is complemented by a molecular system topology, which contains comprehensive information about system composition (i.e., information about all atoms in the molecular system, chemical bonds connecting the atoms and hierarchical organization of atoms into building blocks called residues) and used FF (see the Force Field paragraph below), and output logfiles containing information about system setup, simulation parameters, key parameters of the system such as energy terms at the regular time intervals, etc. The MD simulation thus consists of trajectory file(s), topology file and output logfile(s).

More complicated MD simulation datasets come from specific simulation approaches including, e.g., enhanced sampling techniques. They can involve multiple trajectories (based on the principle of replica-exchange methods, thermodynamic integration, etc.), where set of trajectories (replicas) are run at different conditions defined by so called replica parameter (e.g., temperature or scaling parameter tuning the system’s potential energy) in parallel with regular exchanges (in case of replica-exchange methods) of the replica parameter among the trajectories. Data from replica-exchange simulations can be thus sorted into two complementary sets of simulation paths: (i) continuous trajectories with snapshots following the time evolution of atomic coordinates (each snapshot is evolved from the previous one following equations of motion), and (ii) ensembles containing snapshots following the same value of replica parameter (e.g. temperature or another parameter modifying energy function). Here, for the IDA description and organization, we called datasets created by the simulation software typically in a separated data files as runs (can be either continuous trajectories or ensembles based on the used method) and complementary datasets are called demultiplexed replicas. Note that in case of standard MD simulation, the run, trajectory and ensemble refer to the same single set of snapshots. The set of snapshots of trajectory/run/ensemble between two consecutive exchange attempts will be referred here as chunk (Table 2). Thus, each trajectory/run/ensemble is divided into multiple chunks based on the specified frequency of exchange attempts. Note that in case of standard MD simulation, the chunk span from the beginning to the end of the simulation and thus refers to the same set of snapshots as trajectory, run or ensemble (Figure 3). The most common methods based on replica-exchanges are temperature-replica-exchange MD (T-REMD; multiple replicas are simulated simultaneously at different temperature), Hamiltonian replica exchange (H-REMD, where FF interactions are continuously modified between replicas), a particular implementation of H-REMD called replica exchange with solute tempering (REST or REST2, where only part of molecular system, typically solute atoms are effectively heated up), multi-dimensional REMD (M-REMD), replica exchange with umbrella sampling etc.

**Table 2:** Glossary of key terms along with short description.

| Term | Description |
| --- | --- |
| Simulation | Set of the simulation data consisted of trajectory file(s) containing set of snapshots with atomic coordinates (multiple trajectories in case of replica exchange simulations), topology file with description of a molecular system and FF used, and output log files containing other information about simulation settings |
| Trajectory | Set of snapshots following the time evolution of atomic coordinates, i.e., each snapshot is evolved from the previous one following equations of motion |
| Ensemble | Set of snapshots following the same value of the replica parameter (in case of standard simulation, the ensemble corresponds to the trajectory) |
| Run | Set of snapshots as saved by simulation software into one file. This could be either trajectory or ensemble depending on the used method |
| Demultiplexed replica | Set of snapshots following the complementary paths with respect to runs. |
| Replica parameter | Parameter, which vary among ensembles in the replica exchange simulations. Most typically it is temperature of some scaling parameters modifying energy function. |
| Chunk | Set of snapshots between two consecutive exchange attempts in the replica exchange simulation. All snapshots of the chunk belong to one particular trajectory and one particular ensemble. |

**Figure 3:**
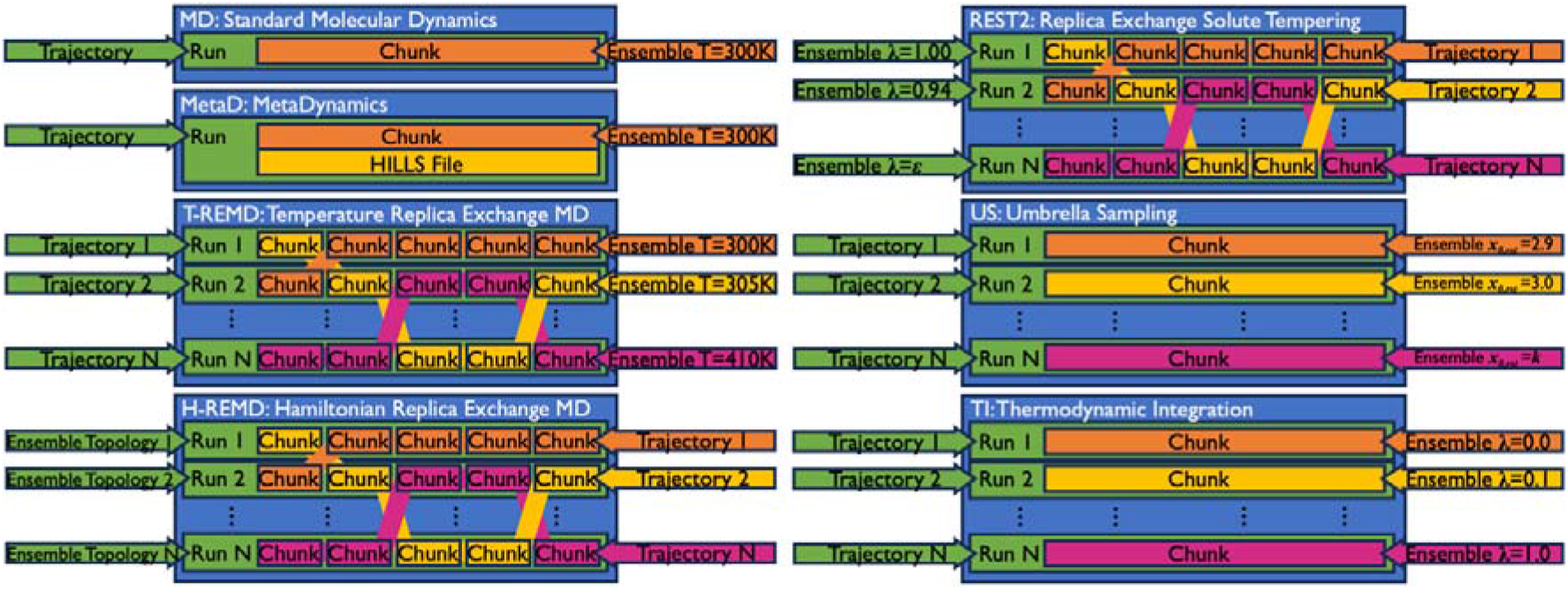
Schematic representation of how IDA data sets can encapsulate simulation data from various MD methods, ensuring seamless interoperability and integration of these data.

Other enhanced sampling approaches are based on the principle of importance sampling, where an artificial biasing force (boost potential) is applied in the simulation to accelerate the sampling in a reduced (low-dimensional) space consisting of selected degrees of freedom denoted as collective variables (CVs). The most common methods are umbrella sampling (US), Gaussian Accelerated MD, Steered MD etc., where external potential is applied to the selected CVs, and metadynamics (MetaD) or its flavor well-tempered variant (WT-MetaD), where history-dependent biasing potential acts on CVs, etc. The simulations using these methods are typically generated by PLUMED plugin for MD simulations (https://www.plumed.org/) and include additional files containing information about bias potential, namely: (i) PLUMED input file with definition of bias(es) and (ii) HILLS file containing the applied biased potential. Combination of both replica-exchange approaches and CV-based methods are also popular, e.g., combination of REST2 with WT-MetaD (ST-MetaD simulations). Some of these methods such as thermodynamic integration (TI) or umbrella sampling (US) use a set of individual trajectories, similarly to the replica exchange methods. The TI simulations involve a series of simulations with different Hamiltonian mixing lambda parameters. Similarly US simulations use different restraint parameters. Thus, in these scenarios, the simulation contains several replicas that are not mutually exchanged. Consequently, each replica has the same structure as a standard MD simulation, meaning that the trajectory, run, or ensemble refer to the same set of snapshots (Figure 3).

In summary, numerous MD approaches have been developed by the community over the past decades, resulting in a diverse set of MD outputs. Here, we show that the innovative IDA structure can incorporate various diverse MD datasets into a standardized format and allow their efficient reusability and interoperability.

### MD parameters

Several key features and MD settings are defined in an input file, which dictates the simulation type, i.e., energy minimization, thermalization, or production MD run. Most common input settings specify output data formats and its saved frequencies, i.e., how often coordinates, velocities, energies, restart files and other outputs are stored. The MD user also specifies constrains and restrains, periodic boundary conditions, treatment of electrostatics, cut-off for non-bonded interactions, barostat (pressure coupling for constant pressure simulations), thermostat (temperature coupling) and the targeted (reference) temperature. The total number of steps is also chosen along with a time step of the integration, e.g., Leap-frog/velocity-Verlet, algorithm. Other settings are rather specific for the particular MD engine and all applied MD settings are saved near the beginning of the output logfile.

### Force Field

FF is a mathematical model used to describe the interactions between atoms and molecules within a molecular system. It comprises a set of equations and parameters that enable enumeration of potential energy and its spatial derivatives for each atom. For instance, AMBER simulation package^16^ uses FF based on the work of Cornell et al.^23^, which approximates the potential energy *V* of molecular system as:

where *k*_*b*_, *k*_*a*_, *V*_*n*_, *r, θ, φ, R*_*ij*_, and *q*_*i*_ stand for spring bond constants, spring angle constants, dihedral amplitude, bond lengths, bond angles, dihedral angles, interatomic distances, and partial atomic charges, respectively.^3^

IDA currently contains non-polarizable molecular mechanics FFs including terms for bonded interactions (e.g., bonds, angles, dihedrals) and non-bonded interactions (e.g., van der Waals forces and electrostatic interactions). This mathematical model is widely used for biomolecular simulations, and here, FF refers to the set of parameters for this particular mathematical model. The IDA database provides separate (and downloadable) datasets with FF parameters for various systems, such as nucleic acids (i.e., RNA, DNA and their building blocks), proteins (also containing parameters for amino acid residues), ligands (typically organic molecules), ions, and solvents. For each molecular type, the database contains multiple FF parameters, such as OL3^24^, DESRES^25^ and ROC^26^ FFs for RNA.

A comprehensive collection of commonly used FFs offers significant benefits to the MD simulation community since many FF parameters are only available upon requests from the original authors. Importantly, the stored FF files contain parameters only for specific molecule types (i.e., RNA) and exclude other (unused and just historically persisting) parameters, e.g., for amino acids, ions, solvents, which is aimed to prevent confusion (mistakes) and make preparation of input files clear and straightforward. Although the IDA currently includes only all-atom non-polarizable molecular mechanics FFs, it is designed to accommodate various FF types. This extensibility includes different flavors of polarizable FFs, coarse-grain FFs, reactive FFs and more.

### Experimental Datasets

The experimental data will be used for validation of MD simulations by comparison with corresponding predictions of measured signals calculated from simulation atomic coordinates, i.e., knowledge extraction data. The experimental data (taken manually from literature) contains information about the measured signals related, e.g., to interaction between particular atoms and thus corresponding to particular distribution of interatomic distance between these atoms, along with the confidence interval (if applicable). Comparing simulation data with experimental observations is particularly straightforward for NMR signals.^27-31^ Consequently, the initial version of IDA implements comparisons exclusively with NMR data. However, it is designed to be extendable to other types of experimental data in the future. Integrating simulation data with experimental datasets significantly enhances their reusability, as validation against experimental benchmarks is crucial for any reevaluation of the simulation data.^3^

### Datasets from Analyses

This part of the database contains data generated by postprocessing of MD simulation, i.e., the knowledge extraction data. Simple analyses produce data about evolution of distance, angle, dihedral, coordination number etc. in time (i.e., for all snapshots) between specific atoms from the simulated system. Thus, in general, the analysis data set contains time development of parameters that is given function of atomic coordinates in each snapshot.

## TECHNICAL VALIDATION

### Analyses

Analyses and comparisons with experimental data are always associated to the particular simulation. They are not directly entered by users but automatically created upon user request. In case of analysis, user select from predefined types of analyses (distance, angle, dihedral, RMSD, etc.) and identify the concerned atoms. The analyses record store the type of the analysis (analysis_type) and its complete definition including mask of concerned atoms in AMBER cpptraj format as a name. Every analysis is associated with set of snapshot_based_on_analysis containing corresponding calculated value (e.g., distance between particular atoms) for each snapshot.

The comparison_with_experimental_data items encompasses the results of comparing simulation data with corresponding experimental data stored in the experimental database. Each simulation is associated via its molecule with set of experimental data. The type of experimental data and experimental conditions such as temperature determine the particular ensemble of the simulation and particular analyses performed in this ensemble to be used for prediction of the corresponding thermodynamic properties that are compared to the experimental values. The comparison_with_experimental_data is thus linked via (i) simulation and molecule to the experimental values and (ii) to the particular analyses of appropriate ensemble.

### Data Curation

Data curation refers to the process of managing, organizing, and maintaining data throughout its lifecycle to ensure its usability, integrity, and long-term preservation. It involves activities such as data collection, validation, annotation, metadata creation, storage, archiving, and dissemination.

The goal of data curation is to enhance the quality, accessibility, and reusability of data for various purposes, including research, analysis, and decision-making. This involves identifying valuable datasets, cleaning, and standardizing data, enriching it with descriptive metadata, and ensuring compliance with relevant policies and standards.

Data stored in the database is machine readable, re-usable, and interoperable. It will be uploaded by an user from manual entries and simulation files (topology, trajectory, and output files). The consistency of the data in these files will be check, e.g., number of atoms as given in topology and trajectory files, number of snapshots in all trajectory files and output logfiles etc. The data manually uploaded by the user are highlighted in Figure 2. Other data will be parsed from the files uploaded by the user and during their processing they will be checked. Notably, the key annotation of data such as assignment of the FF to the simulation will be done automatically during processing the input files, which will significantly improve integrity of the data. Storing the simulation data in the internal database structure via parsing of primary files produced by MD simulations is a flexible choice enabling parsing of various file formats including, e.g., various software, future format updates etc. Once the respective items are uploaded, they will be locked and subsequently assigned persistent identifiers (PIDs), such as those based on the EUDAT-B2HANDLE service (https://eudat.eu/service-catalogue/b2handle).

### Metadata Extraction and FAIR Data Aspects

Metadata encompass all essential information related to the research data (*cf*. Figure 2). Extracting metadata from the database involves running a database query. Users can extract metadata pertaining to molecules, simulations, FFs, and experimental data. The extracted metadata will include the following mandatory details:

⍰ Identifier: PID assigned to the respective item (as mentioned above; in case of data(sets) with elevated importance, we consider additionally assigning a DOI – a Digital Object Identifier).
⍰ Creator Name: The individual or entity responsible for creating the data.
⍰ Publication Date: The date when the data was published or made available.
⍰ Title: The title or name associated with the data.
⍰ Description: A brief overview or description of the data.
⍰ Rights: Information regarding the usage rights or restrictions associated with the data.
⍰ Additional information specific to each database section: This may include details relevant to the specific category of data, such as simulation parameters, experimental conditions, or molecular properties.

The metadata format is standardized across the database, with metadata extracted in English and structured according to the mandatory tagged fields in https://zenodo.org/records/7040047 using XML. Creator and uploader information will be stored in a separate table, with associations made through unique identifiers, such as email addresses. This set of metadata has been designed to form a superset of the mandatory fields in the DataCite 4.5 metadata standard (https://schema.datacite.org). Along with the assignment of PIDs, adhering to such a widely accepted, cross-disciplinary metadata standard is crucial for ensuring that the IDA complies with the FAIR principles of research data management. Additionally, this compliance enables us to mint DOIs (via DataCite) for selected datasets.

## Supporting information

Supporting Information to the article.

## USAGE NOTES

In this work, we described the IDA database (Integrated Database of force fields (FFs) and datasets from experiments and MD simulations), an innovative scheme addressing the primary challenges in the development and utilization of databases for MD simulations. IDA’s adherence to FAIR principles should promote transparency and facilitate collaboration across research communities. The database’s compliance with these principles, coupled with its web interface accessibility, underscores its potential to become a cornerstone resource in the MD simulation community.

IDA is designed to answer main challenges in storing and efficient data reusability, i.e., volume of output data, diverse formats and structures of datasets generated by different MD engines and approaches. IDA addresses these tasks by providing a standardizable format that merges, classifies, and stores datasets, facilitating efficient reuse and data mining from the vast amounts of data generated by diverse approaches and stored together. This capability allows for valuable insights into the performance and accuracy of FFs and the effects of specific simulation settings, which are currently underutilized in existing databases.

IDA is also allowing integration of MD data with other relevant data sources, including FF parameters and experimental data. By interconnecting these datasets, IDA enables the assessment of the relevance of MD simulations, testing of new simulation methods, and development of advanced analytics tools. This integration is crucial for enhancing the reliability and predictive power of MD simulations, particularly when coupled with machine learning (ML) tools that require accurate, consistent, and well-structured data. The seamless integration promoted by IDA supports innovation in fields such as drug discovery, materials science, and biotechnology.

In conclusion, IDA represents a significant advancement in the management and utilization of MD simulation data. By addressing key challenges related to data volume, diversity, and integration, IDA fosters a more robust, interoperable, and reusable data ecosystem, paving the way for enhanced scientific discoveries and innovations.

## CODE AVAILABILITY

IDA database is available via GUI at https://ida.4sims.eu/simulations.

## ACKNOWLEDGEMENTS

This research received the support of EXA4MIND, a European Union′s Horizon Europe Research and Innovation programme under grant agreement N° 101092944. Views and opinions expressed are however those of the author(s) only and do not necessarily reflect those of the European Union or the European Commission. Neither the European Union nor the granting authority can be held responsible for them. This work was also supported by the Czech Science Foundation to V.M. and J.S. (grant number 23-05639S). M.O. and P.B. were supported by ERDF/ESF project TECHSCALE (No. CZ.02.01.01/00/22_008/0004587).

## AUTHOR CONTRIBUTIONS

P.B., V.M., J.M., and M.O. conceived the project, V.M. performed the calculations, D.C., R.F., N.P., V.M., and V.P. curated the data and implemented the database. P.B., V.M., and M.O. wrote the manuscript with contributions from all authors.

## SUPPLEMENTARY INFORMATION

Supplementary Information of IDA: Integrated Database of Force-Field Parameters, Experimental Measurements and Molecular Dynamics Simulations.

## REFERENCES

(1) Cheatham, T. E., 3rd; Case, D.A. Twenty-five years of nucleic acid simulations. Biopolymers 2013, 99 (12), 969–977. DOI: 10.1002/bip.22331.

(2) Smith, L. G.; Zhao, J.; Mathews, D. H.; Turner, D. H. Physics-based all-atom modeling of RNA energetics and structure. Wiley Interdiscip Rev RNA 2017, 8 (5). DOI: 10.1002/wrna.1422.

(3) Sponer, J.; Bussi, G.; Krepl, M.; Banas, P.; Bottaro, S.; Cunha, R. A.; Gil-Ley, A.; Pinamonti, G.; Poblete, S.; Jurecka, P.; et al. RNA Structural Dynamics As Captured by Molecular Simulations: A Comprehensive Overview. Chem Rev 2018, 118 (8), 4177–4338. DOI: 10.1021/acs.chemrev.7b00427.

(4) Paloncyova, M.; Pykal, M.; Kuhrova, P.; Banas, P.; Sponer, J.; Otyepka, M. Computer Aided Development of Nucleic Acid Applications in Nanotechnologies. Small 2022, 18 (49), e2204408. DOI: 10.1002/smll.202204408 From NLM Medline.

(5) Berman, H. M.; Westbrook, J.; Feng, Z.; Gilliland, G.; Bhat, T. N.; Weissig, H.; Shindyalov, I. N.; Bourne, P. E. The Protein Data Bank. Nucleic Acids Res 2000, 28 (1), 235–242. DOI: 10.1093/nar/28.1.235 From NLM Medline.

(6) van der Kamp, M. W.; Schaeffer, R. D.; Jonsson, A. L.; Scouras, A. D.; Simms, A. M.; Toofanny, R. D.; Benson, N. C.; Anderson, P. C.; Merkley, E. D.; Rysavy, S.; et al. Dynameomics: a comprehensive database of protein dynamics. Structure 2010, 18 (4), 423–435. DOI: 10.1016/j.str.2010.01.012 From NLM Medline.

(7) Meyer, T.; D’Abramo, M.; Hospital, A.; Rueda, M.; Ferrer-Costa, C.; Perez, A.; Carrillo, O.; Camps, J.; Fenollosa, C.; Repchevsky, D.; et al. MoDEL (Molecular Dynamics Extended Library): a database of atomistic molecular dynamics trajectories. Structure 2010, 18 (11), 1399–1409. DOI: 10.1016/j.str.2010.07.013 From NLM Medline.

(8) Hensen, U.; Meyer, T.; Haas, J.; Rex, R.; Vriend, G.; Grubmuller, H. Exploring protein dynamics space: the dynasome as the missing link between protein structure and function. PLoS One 2012, 7 (5), e33931. DOI: 10.1371/journal.pone.0033931 From NLM Medline.

(9) Thibault, J. C.; Facelli, J. C.; Cheatham, T. E., 3rd. iBIOMES: managing and sharing biomolecular simulation data in a distributed environment. J Chem Inf Model 2013, 53 (3), 726–736. DOI: 10.1021/ci300524j From NLM Medline.

(10) Hospital, A.; Andrio, P.; Cugnasco, C.; Codo, L.; Becerra, Y.; Dans, P. D.; Battistini, F.; Torres, J.; Goni, R.; Orozco, M.; et al. BIGNASim: a NoSQL database structure and analysis portal for nucleic acids simulation data. Nucleic Acids Res 2016, 44 (D1), D272–278. DOI: 10.1093/nar/gkv1301 From NLM Medline.

(11) Beltran, D.; Hospital, A.; Gelpi, J. L.; Orozco, M. A new paradigm for molecular dynamics databases: the COVID-19 database, the legacy of a titanic community effort. Nucleic Acids Res 2024, 52 (D1), D393–D403. DOI: 10.1093/nar/gkad991 From NLM Medline.

(12) Hospital, A.; Orozco, M. MD-DATA: the legacy of the ABC Consortium. Biophysical Reviews 2024, 1–3.

(13) Kiirikki, A. M.; Antila, H. S.; Bort, L. S.; Buslaev, P.; Favela-Rosales, F.; Ferreira, T. M.; Fuchs, P. F. J.; Garcia-Fandino, R.; Gushchin, I.; Kav, B.; et al. Overlay databank unlocks data-driven analyses of biomolecules for all. Nat Commun 2024, 15 (1), 1136. DOI: 10.1038/s41467-024-45189-z From NLM Medline.

(14) Antila, H. S.; Kav, B.; Miettinen, M. S.; Martinez-Seara, H.; Jungwirth, P.; Ollila, O. S. Emerging era of biomolecular membrane simulations: automated physically-justified force field development and quality-evaluated databanks. The Journal of Physical Chemistry B 2022, 126 (23), 4169–4183.

(15) Wilkinson, M. D.; Dumontier, M.; Aalbersberg, I. J.; Appleton, G.; Axton, M.; Baak, A.; Blomberg, N.; Boiten, J. W.; da Silva Santos, L.B.; Bourne, P. E.; et al. The FAIR Guiding Principles for scientific data management and stewardship. Sci Data 2016, 3, 160018. DOI: 10.1038/sdata.2016.18 From NLM Medline.

(16) Case, D. A.; Cheatham, T. E.; Darden, T.; Gohlke, H.; Luo, R.; Merz, K. M.; Onufriev, A.; Simmerling, C.; Wang, B.; Woods, R. J. The Amber biomolecular simulation programs. Journal of computational chemistry 2005, 26 (16), 1668–1688.

(17) Gutmanas, A.; Adams, P. D.; Bardiaux, B.; Berman, H. M.; Case, D. A.; Fogh, R. H.; Guntert, P.; Hendrickx, P. M.; Herrmann, T.; Kleywegt, G. J.; et al. NMR Exchange Format: a unified and open standard for representation of NMR restraint data. Nat Struct Mol Biol 2015, 22 (6), 433–434. DOI: 10.1038/nsmb.3041 From NLM Medline.

(18) Ulrich, E. L.; Baskaran, K.; Dashti, H.; Ioannidis, Y. E.; Livny, M.; Romero, P. R.; Maziuk, D.; Wedell, J. R.; Yao, H.; Eghbalnia, H. R.; et al. NMR-STAR: comprehensive ontology for representing, archiving and exchanging data from nuclear magnetic resonance spectroscopic experiments. J Biomol NMR 2019, 73 (1-2), 5–9. DOI: 10.1007/s10858-018-0220-3 From NLM Medline.

(19) Roe, D. R.; Cheatham, T. E. PTRAJ and CPPTRAJ: Software for Processing and Analysis of Molecular Dynamics Trajectory Data. Journal of Chemical Theory and Computation 2013, 9 (7), 3084–3095. DOI: 10.1021/ct400341p.

(20) Abraham, M. J.; Murtola, T.; Schulz, R.; Páll, S.; Smith, J. C.; Hess, B.; Lindahl, E. GROMACS: High performance molecular simulations through multi-level parallelism from laptops to supercomputers. SoftwareX 2015, 1, 19–25.

(21) Eastman, P.; Swails, J.; Chodera, J. D.; McGibbon, R. T.; Zhao, Y.; Beauchamp, K. A.; Wang, L. P.; Simmonett, A. C.; Harrigan, M. P.; Stern, C. D.; et al. OpenMM 7: Rapid development of high performance algorithms for molecular dynamics. PLoS Comput Biol 2017, 13 (7), e1005659. DOI: 10.1371/journal.pcbi.1005659 From NLM Medline.

(22) Adjoua, O.; Lagardere, L.; Jolly, L. H.; Durocher, A.; Very, T.; Dupays, I.; Wang, Z.; Inizan, T. J.; Celerse, F.; Ren, P.; et al. Tinker-HP: Accelerating Molecular Dynamics Simulations of Large Complex Systems with Advanced Point Dipole Polarizable Force Fields Using GPUs and Multi-GPU Systems. J Chem Theory Comput 2021, 17 (4), 2034–2053. DOI: 10.1021/acs.jctc.0c01164 From NLM PubMed-not-MEDLINE.

(23) Cornell, W. D.; Cieplak, P.; Bayly, C. I.; Gould, I. R.; Merz, K. M.; Ferguson, D. M.; Spellmeyer, D. C.; Fox, T.; Caldwell, J. W.; Kollman, P. A. A second generation force field for the simulation of proteins, nucleic acids, and organic molecules (vol 117, pg 5179, 1995). J Am Chem Soc 1996, 118 (9), 2309–2309.

(24) Zgarbova, M.; Otyepka, M.; Sponer, J.; Mladek, A.; Banas, P.; Cheatham, T. E.; Jurecka, P. Refinement of the Cornell et al. Nucleic Acids Force Field Based on Reference Quantum Chemical Calculations of Glycosidic Torsion Profiles. J Chem Theory Comput 2011, 7 (9), 2886–2902. DOI: 10.1021/ct200162x.

(25) Tan, D.; Piana, S.; Dirks, R. M.; Shaw, D. E. RNA force field with accuracy comparable to state-of-the-art protein force fields. Proc Natl Acad Sci U S A 2018, 115 (7), E1346–E1355. DOI: 10.1073/pnas.1713027115.

(26) Aytenfisu, A. H.; Spasic, A.; Grossfield, A.; Stern, H. A.; Mathews, D. H. Revised RNA Dihedral Parameters for the Amber Force Field Improve RNA Molecular Dynamics. J Chem Theory Comput 2017, 13 (2), 900–915.

(27) Condon, D. E.; Kennedy, S. D.; Mort, B. C.; Kierzek, R.; Yildirim, I.; Turner, D. H. Stacking in RNA: NMR of Four Tetramers Benchmark Molecular Dynamics. J Chem Theory Comput 2015, 11 (6), 2729–2742. DOI: 10.1021/ct501025q.

(28) Bottaro, S.; Bussi, G.; Kennedy, S. D.; Turner, D. H.; Lindorff-Larsen, K. Conformational ensembles of RNA oligonucleotides from integrating NMR and molecular simulations. Sci Adv 2018, 4 (5), eaar8521. DOI: 10.1126/sciadv.aar8521.

(29) Zhao, J.; Kennedy, S. D.; Berger, K. D.; Turner, D. H. Nuclear Magnetic Resonance of Single-Stranded RNAs and DNAs of CAAU and UCAAUC as Benchmarks for Molecular Dynamics Simulations. J Chem Theory Comput 2020, 16 (3), 1968–1984. DOI: 10.1021/acs.jctc.9b00912.

(30) Bottaro, S.; Nichols, P. J.; Vogeli, B.; Parrinello, M.; Lindorff-Larsen, K. Integrating NMR and simulations reveals motions in the UUCG tetraloop. Nucleic Acids Res 2020, 48 (11), 5839–5848. DOI: 10.1093/nar/gkaa399.

(31) Zhao, J.; Kennedy, S. D.; Turner, D. H. Nuclear Magnetic Resonance Spectra and AMBER OL3 and ROC-RNA Simulations of UCUCGU Reveal Force Field Strengths and Weaknesses for Single-Stranded RNA. J Chem Theory Comput 2022, 18 (2), 1241–1254. DOI: 10.1021/acs.jctc.1c00643.

