## Supporting Information to the article. for "Integrated Database of Force-Field Parameters, Experimental Measurements and Molecular Dynamics Simulations"

**by**

*Pavel Banáš, Vojtěch Mlýnský, David Číž, Radek Furmánek, Nestor Pilat, Viktoria Pauw, Stephan Hachinger, Jiří Šponer, Jan Martinovič, and Michal Otyepka*

### Supplementary Tables

Supplementary tables provide detailed information for each section of the IDA database (see Figure 2 in the maintext). Each attribute is expanded with an example value, a data type hint, and a brief description explaining its significance.

**Table S1:** Details about **molecule** section.

**molecule**

| Attribute | Value | Type | Description |
|---|---|---|---|
| Id | 88 | bigint | ID of item |
| molecule_name | r(UUUU) | string | Name of the molecule in the database. |
| molecule_sequence | 5’-CGCGAATTCGGC-3’ | string | Sequence of respective molecule, either, nucleic acid or protein, from 5’ to 3’ end or from N- to C-terminus, for nucleic acid or protein, respectively. |
| molecule_type | DNA, RNA, DNA/RNA | array | Type of molecule, DNA, RNA, DNA/RNA, protein ... |
| molecule_description | r(UUUU) tetranucleotide | string | Brief description of the molecule. |
| molecule_organism | artificial/none/E.coli | enum | Source organism. |
| molecule_organism_desc | “artificial based on…” | string | Additional description of the respective molecule. |
| data_upload_timestamp | 14.1.2024 | datetime | Time stamp of molecule upload into database. |
| upload_user_id | 456975 | bigint | Identification of uploading. |

**Table S2:** Details about **simulation** section.

| **simulation** | | | |
|---|---|---|---|
| Attribute | Value | Type | Description |
| Id | 123456 | bigint | ID of simulation |
| DOI | 10.1234/sim_UUUUU123456 | string | DOI assigned to simulation item after upload into database, after DOI assignment the item is locked and cannot be further modified. |
| author_name | "Rudolf" | varchar | Name of the author, who created/executed the MD simulation. |
| upload_user_id | 456975 | bigint | Identification of the user. |
| software_name | "PMEMD" | varchar | Name of the simulation software. |
| software_settings | "64 float accuracy, …" | string | Setting(s) used for execution of MD simulation. |
| software_version | 1.2.15 - cpu | varchar | Version of the software used. |
| simulation_creation_timestamp | 2021-07-21T11:30:18.0 | datetime | Date of simulation creation parsed from log file. |
| simulation_upload_timestamp | 2024-03-27T18:34:38.24 | datetime | Time stamp of simulation upload into database. |
| simulation_description | "extending work of [cite], exploring the effect of …" | string | Additional information about respective MD simulation. |
| sim_name | "uuuu-ROC-TIP3P-0.1NaCl" | varchar | Author specified name of the simulation. |
| solvent | "WAT, MET, ACE" | array | Solvent types taken from topology file. |
| solvent_ff | OPC | string | Name associated with the solvent ff parameters. |
| ion | "Cl-, Ca2+, K+" | array | Ion types taken from topology file. |
| ion_ff | Joung-Cheatham | string | Name associated with ion ff parameters. |
| ionic_concentration (mM) | 150 | float | Estimation of ionic concentration. |
| n_snapshots | 10000 | bigint | Number of snapshots. |
| stripping_mask | :WAT,:20-30, None | string | Stripping mask. |
| ensemble_parameter_type | temperature OR lambda etc. | varchar | Temperature or topology scaling parameter in replica exchange simulations. |
| time_step (ps) | 0.004 | float | Integration time step in ps. |
| total_simulation_time (ns) | 10000 | Float | Total simulation time. |
| thermostat | "Nose-Hoover", "Langevin", … | String | Used simulation thermostat. |
| barostat | "Berensen", … | String | Used simulation barostat. |
| ensemble_ladder | [300 K, 310 K], [λ0.8, λ0.9] | array | Complete list of temperatures or topology scalling parameters in replica exchange simulations. |
| restraint_file | "restraints.dat" or NONE | blob | Additional input file(s) with external force-field modifications. |
| molecule_id | 88 | bigint(fk) | Molecule identifier. |
| ensemble | NPT or NVT | varchar | Used simulation ensemble, e.g., [N,p,T] |
| temperature | 300 K or NONE (T-REMD) | float | Simulation temperature. If temperature is replica parameter, e.g. in T-REMD, the temperature here is set to NONE and temperatures of ensemble paths are specified in the particular path records and summarized here as ensemble_ladder. |
| pressure | 1 atm or NONE (e.g. NVT) | float | Used simulation pressure. |

**Table S3:** Details about **path** section.

| **path** | | | |
|---|---|---|---|
| Attribute | Value | Type | Description |
| id | 654000 | bigint | Unique identifier. |
| traj_id | 1, 2, or -1 for NONE | int(fk) | Trajectory identifier (if path corresponds to the trajectory). |
| ensemble_param | 300K, 0.9, or -1.0 for NONE | float | Ensemble parameter, i.e. temperature/scalling parameter value (if path corresponds to the ensemble). |
| simulation_id | 123456 | bigint(fk) | Simulation identifier. |
| plumed | plumed.dat or NONE | blob | Additional input file with biases (metadynamics simulations) and/or external force-field modifications. |

**Table S4:** Details about **chunk** section.

| **chunk** | | | |
|---|---|---|---|
| Attribute | Values | Type | Description |
| id | 654321 | bigint | Unique identifier. |
| path_traj_id | 654000 | bigint(fk) | Index to path identifier of the trajectory containing this chunk. |
| path_ensemble_id | 654321 | bigint | Index to path identifier of the ensemble containing this chunk. |
| order_for_path | 45/10000 | int | Possition of the chunk in the its trajectory and ensemble |
| chunk_size | 2 | int | Number of snapshots in the chunk. |
| traj_id | 1 OR 2 OR 3 … | int | Index of chunk's trajectory |
| ensemble_param | 300(K) OR 0.8 (0.9) | float | Replica parameter, i.e., temperature/scalling parameter value, of the chunk's ensemble. |

**Table S5:** Details about **snapshot** section.

| **snapshot** | | | |
|---|---|---|---|
| Attribute | Values | Type | Description |
| id | 111111 | bigint | Snapshot identifier. |
| time | 0.12 | float(ps) | Simulation time corresponding to the snapshot |
| chunk_id | 654321 | bigint(fk) | Chunk identifier. |
| box | [0.1,0.4,0.55] | array | Size of the solvent box in this snapshot |

**Table S6:** Details about **coordinates** section.

**coordinates**

| Attribute | Values | Type | Description |
|---|---|---|---|
| X | 954.1 | Float | First Cartesian coordinate defying position of the point (atom) in three-dimensional space. |
| Y | 12.44 | Float | Second Cartesian coordinate defying position of the point (atom) in three-dimensional space. |
| Z | 47 | Float | Third Cartesian coordinate defying position of the point (atom) in three-dimensional space. |
| snapshot_id | 111111 | BigInt(fk) | Snapshot identifier. |
| atom_id | 999999 | BigInt(fk) | Atom identifier. |

**Table S7:** Details about **atoms** section.

**atoms**

| Attribute | Values | Type | Description |
|---|---|---|---|
| id | 999999 | bigint | Atom identifier. |
| simulation_id | 123456 | bigint(fk) | Simulation identifier. |
| element | C OR N OR H | String | Atomic element. |
| name | C4 | String | Atom name. |
| resname | Gly | String | Residue name. |
| resid | 10 | int | Residue identifier. |

**Table S8:** Details about **analysis** section.

**analysis**

| Attribute | Value | Type | Description |
|---|---|---|---|
| id | 123456 | bigint | Analysis identifier. |
| path_id | 654321 | Integer | Path identifier |
| parameter_type | distance, dihedral, | String | Type of parameter being analyzed, e.g., distance, angle, etc. |
| name | dist_:2@C5'_:2@C4' | String | The name given to the parameter for identification purposes. |

**Table S9:** Details about **snapshot_based_on_analysis** section.

**snapshot_based_on_analysis**

| Attribute | Value | Type | Description |
|---|---|---|---|
| id | 777777 | bigint | A unique identifier |
| snapshot_id | 65412 | bigint(fk) | The snapshot identifier |
| analysis_id | 456123 | bigint(fk) | Analysis identifier. |
| computed_value | [41.412] | array | Calculated value of particular analysis in the particular snapshot. |

**Table S10:** Details about **comparison_with_experimental_data** section.

**comparison_with_experimental_data**

| Attribute | Values | Data | Description |
|---|---|---|---|
| id | 777777 | bigint | Unique identifier |
| simulation_id | 456123 | bigint(fk) | The simulation identifier |
| ff_id | 456123 | bigint(fk) | Force field identifier |
| value | 3.2 | float | Quantitative measure obtained from the comparison of simulation with experimental data. |
| significance | 1.2 | float | Indicates the statistical significance of the value measuring the agreement of simulation data with experimental values using reweighting. |

**Table S11:** Details about **FF** section.

**FF**

| Attribute | Values | Type | Description |
|---|---|---|---|
| Id | 654321 | bigint | Unique identifier for each force field entry. |
| DOI | 10.1234/FFOL3 | string | Digital Object Identifier for referencing the force field. |
| FF_name | "Chen-Garcia", "OL3" | varchar | Names associated with the force field. |
| molecule_type | DNA, RNA | array | Types of molecules that the force field is applicable to. |
| FF_library_file | RNA.OL3.lib | text | File containing the library of the force field parameters. |
| FF_parm_file | parm.RNA.OL3.dat | text | File containing parameter data for the force field. |
| FF_leaprc_file | leaprc.RNA.OL3 | text | File containing input data for parameter and topology files generation in AMBER. |
| data_publication_time stamp | 43844 | datetime | Date when the data related to this force field was published. |
| data_upload_timestam p | 45305 | datetime | Date when this specific data entry was uploaded into the database. |
| reference_article_DOI | 10.1021/ct200162x | string | DOI of the reference article where this force field is described or used. |
| author_name | Zgarbova | varchar | Name of one of the authors associated with this force field or its reference article. |
| upload_user_id | 456975 | varchar | Assigned identifier to user who uploaded the data. |

**Table S12:** Details about **experimental data** section.

**experimental data**

| Attribute | Values | Type | Description |
|---|---|---|---|
| Id | 654321 | bigint | Unique identifier for each entry in the experimental data. |
| DOI | 10.1234/exp_UUCG1234 | string | Digital Object Identifier, a unique alphanumeric string assigned to this database item. |
| molecule_id | 88 | bigint(fk) | Foreign key reference to a specific molecule in the database. |
| exp_data_type | NMR-NOE, NMR-3J | string | Types of experimental data obtained, indicating specific NMR methods used. |
| values | exp_data.dat | blob | Binary large object containing the raw experimental data values. |
| pdb_id | 2KOC | text | Four letter code of the molecule stored at PDB structural database (https://www.rcsb.org/). |
| pdb_doi | https://doi.org/10.2210/pdb2KOC/pdb | text | Digital identifier of the molecule at PDB structural database (https://www.rcsb.org/). |
| data_publication_timestamp | 2021-07-21T11:30:18.0 | datetime | The date and time when the experimental data was published publicly. |
| data_upload_timestamp | 2022-04-11T14:40:11.0 | datetime | The date and time when the experimental data was uploaded to the database. |
| reference_article_DOI | 10.1093/nar/gkp956 | string | Digital Object Identifier of the reference article associated with this dataset. |
| author_name | "Harald Schwalbe" | string | Name of the author who conducted or is associated with this experiment. |
| upload_user_id | 456975 | string | Assigned identifier to the user who uploaded this dataset to the database. |
| temperature (K) | 275 | float | Temperature at which this particular set of experimental data was collected. |
